# Social behaviour is a key determinant of virome diversity in New Zealand passerines

**DOI:** 10.64898/2026.10.15.759493

**Authors:** Kristina M Hames, Stephanie J Waller, Rebecca K French, Janelle R Wierenga, Benjamin Perry, Fiona Robertson, Bruce C. Robertson, Jemma L Geoghegan

**Affiliations:** Department of Microbiology and Immunology, University of Otago, Dunedin, Aotearoa New Zealand; Department of Zoology, University of Otago, Dunedin, Aotearoa New Zealand; The Otago Peninsula Eco Restoration Alliance, Portobello, Dunedin, Aotearoa New Zealand; New Zealand Institute for Bioeconomy Science Limited, Invermay Agricultural Centre, Dunedin, Aotearoa New Zealand

**Author notes:** Author for correspondence: Jemma L Geoghegan.

**Keywords:** virome, viral ecology, metatranscriptomics, passerines, environmental surveillance, New Zealand

## Abstract

Wild birds are important reservoirs of viral diversity and can play key roles in the emergence and dispersal of infectious diseases. Their ability to harbour multiple infections without overt disease, combined with their mobility and ecological interactions, creates opportunities for viruses to spread across species and geographic regions. Despite this, the factors shaping avian viromes remain poorly understood, particularly in Aotearoa New Zealand, where endemic and introduced bird species coexist within a relatively isolated ecosystem. This study used metatranscriptomic sequencing to characterise the viromes of 443 endemic and introduced passerines sampled over four years within a shared ecological setting. A total of 217 viruses, representing 69 viral species across 14 families, were identified, including 54 previously undescribed viruses. Many were closely related to viruses reported from birds in Asia, North America, Europe and Australia, suggesting historical or ongoing viral exchange through migration, co-divergence or cross-species transmission. Introduced passerines harboured greater viral diversity than endemic species, while social behaviour was a key predictor of virome composition with flocking species exhibiting higher viral richness and diversity than solitary or pair-living taxa. In contrast, non-viral microbial communities were primarily shaped by host species and diet, indicating that viral communities are influenced by ecological processes distinct from those associated with the non-viral microbiome. These findings provide new insight into the ecological and behavioural drivers of avian viral diversity.

## Introduction

Wild birds are important reservoirs of viral diversity, harbouring a wide range of viruses with potential relevance to wildlife, livestock and human health (Nabi et al., 2021). Many viral infections in wild birds are asymptomatic, although this varies among host species and viruses (Chan et al., 2015). Combined with the high mobility of some species, these infections may facilitate viral maintenance, dispersal and evolution across broad spatial and taxonomic scales (Wille & Holmes, 2020). Among avian hosts, passerines (order Passeriformes) are particularly notable due to their diversity, abundance and ecological ubiquity, comprising more than half of all bird species globally (Williams et al., 2023). Despite their ecological prominence, the diversity and ecological drivers of passerine viral communities remain relatively understudied.

Viral transmission within bird communities is strongly influenced by ecological and behavioural factors (Borremans et al., 2019). Shared feeding sites, water sources and roosting areas create opportunities for transmission within and between species, while social behaviours such as flocking increase contact rates and may facilitate both viral maintenance and transmission (Farine et al., 2014). These processes can be amplified in human-modified environments (Sumasgutner et al., 2023). Understanding how host ecology shapes viral diversity is therefore central to understanding patterns of viral transmission and emergence in wild bird populations.

Aotearoa New Zealand provides a unique environment for studying avian viral ecology. Its long geographic isolation has resulted in a highly distinctive avifauna, with approximately half of all bird species being endemic (Maderspacher, 2022; Wallis & Trewick, 2009). Following human settlement, approximately 120 non-native bird species were introduced, of which around 40 established self-sustaining wild populations (Pipek et al., 2019). These introduced species now coexist with native birds, particularly within urban and agricultural landscapes. Such novel ecological assemblages create opportunities for pathogen exchange between endemic and introduced hosts (Campbell et al., 2016), yet the extent to which host ecology influences viral diversity and sharing within these communities remains largely unknown.

Previous studies have revealed diverse viral communities in New Zealand birds, including endemic and introduced passerines from the South Island (French, Filion, et al., 2022), demonstrating that these hosts harbour substantial viral diversity. However, this earlier work involved a comparatively small sample and provided a single temporal snapshot. Consequently, the ecological factors shaping avian virome composition, and the consistency of these patterns over time, remain poorly resolved. Understanding how host traits such as social behaviour, diet and taxonomy influence viral diversity is important for identifying the mechanisms that facilitate viral transmission. This is particularly relevant in human-modified environments, which may act as transmission interfaces where viruses circulate among phylogenetically distinct species and potentially emerge in new host populations (Ha et al., 2013).

Here, we applied metatranscriptomic sequencing to characterise viral communities in 443 passerines sampled over four years from a mixed assemblage of endemic and introduced species in Dunedin, New Zealand. Using the same sequencing data we also characterised broader host-associated non-viral microbial communities. We tested whether host taxonomy, diet, social behaviour and endemic status influence virome composition and patterns of viral sharing, and compared these drivers with those shaping the broader non-viral microbial communities.

## Methods and materials

### Ethics

This work was approved by the University of Otago’s Animal Ethics Committee (AUP-21-84) and the New Zealand Department of Conservation (Authorisation Number: 94794-FA). This project was endorsed by the Ngāi Tahu Research Consultation Committee at the University of Otago.

### Study location and bird species composition

Sampling took place within a single backyard in the small residential area of Dunedin, New Zealand (Figure 1). The site featured a mix of garden trees and surrounding vegetation, as well as shared food and water sources used by multiple bird species, creating opportunities for interspecific contact and potential virus transmission. The site attracted a diverse community of passerine birds, including endemic species such as the bellbird (*Anthornis melanura*), the tūī (*Prosthemadera novaeseelandiae*) and the self-introduced silvereye/tauhou (*Zosterops lateralis*), which naturally colonised New Zealand from Australia. The site also supported a range of introduced passerines including house sparrow (*Passer domesticus*), dunnock (*Prunella modularis*), blackbird (*Turdus merula*), song thrush (*Turdus philomelos*), greenfinch (*Chloris chloris*), goldfinch (*Carduelis carduelis*), chaffinch (*Fringilla coelebs*), and starling (*Sturnus vulgaris*).

**Figure 1.**
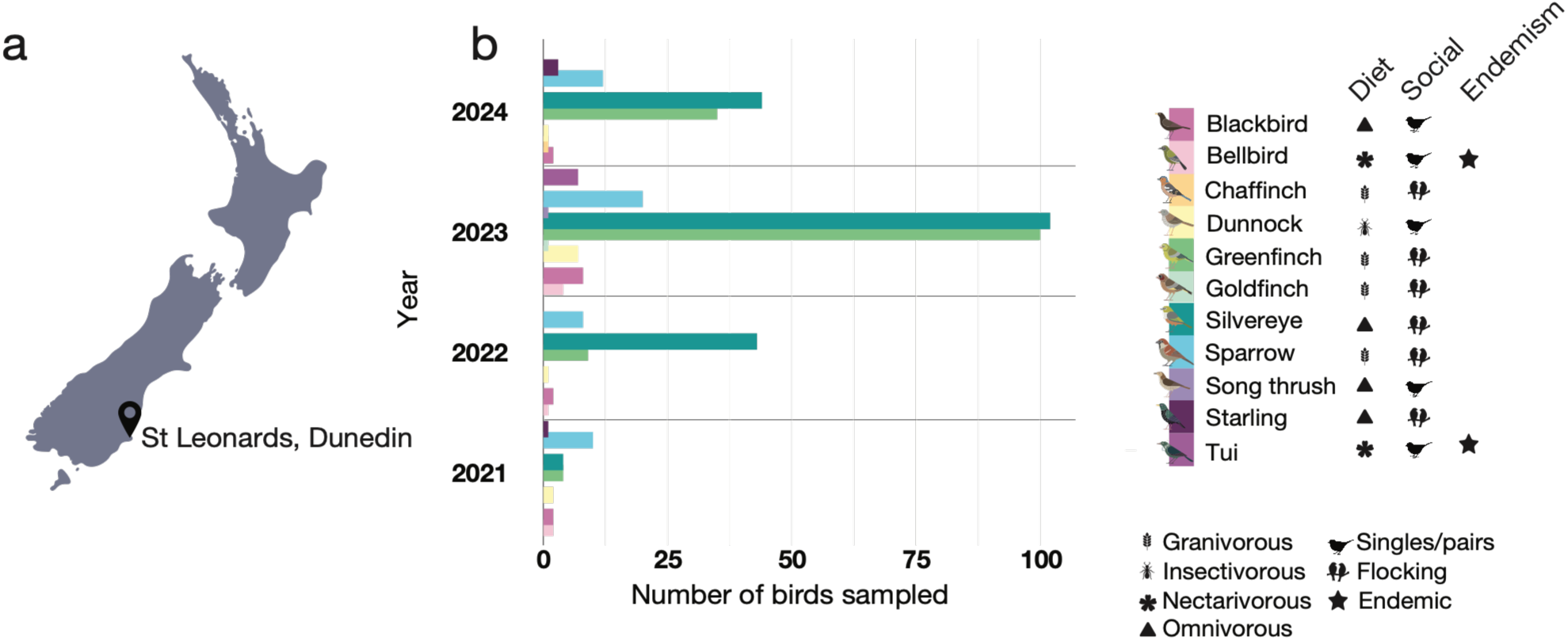
**(a)** Map of New Zealand indicating the study site in St Leonards, Dunedin (South Island of New Zealand). **(b)** The number of individuals captured for each species throughout the study period. Accompanying icons denote endemic status, predominant diet and social behaviour for each species.

### Sample collection

Fieldwork was conducted between the 24 November 2021 and 14 November 2024. A total of 443 birds, representing 11 passerine species, were captured using low-canopy mist nets (Supplementary Table S1). Silvereyes were additionally captured using a drop-lid trap, as their behaviour and use of the feed area made this method more effective for this species. Sugar water and bird feed were used to attract the birds into the backyard. Upon capture, each bird was deemed healthy, measured and weighed, and a cloacal swab was collected using a sterile flock swab (Puritan). The entire tip of the swab was gently inserted into the cloaca and gently rotated multiple times. The swab tip was then cut using scissors sterilised with 70% ethanol and placed into a sterile 2 mL collection tube containing 1mL of DNA/RNA Shield (Zymo Research). Samples were stored at −20°C immediately after collection, then transferred to −80°C until processing. All procedures followed ethical guidelines and relevant permits to ensure animal welfare.

### RNA extraction and sequencing

Total RNA was extracted following the ZymoBIOMICS MagBead RNA kit (Zymo Research). The frozen cloacal swabs were thawed and transferred using sterile forceps into individual ZR BashingBead lysis tubes (0.1 and 0.5 mm) containing 750 µl of fresh DNA/RNA Shield (Zymo Research). The samples were homogenised for five minutes using a bead beater (BioSpec Products Inc.) disruptor. The extracted RNA was eluted into 50μL DNase/RNase-free water and RNA concentrations were quantified using a NanoDrop Spectrophotometer (ThermoFisher). Equal quantities of RNA from individual birds were pooled into 52 groups by species and year caught (2021-2024). To manage instances with a larger number of individuals per species and year, these were divided into multiple pools, with each library containing RNA from up to 20 individuals (Supplementary Figure S1). RNA sequencing was performed on the Illumina NovaSeqX platform after ribosomal RNA removal with Ribo-Zero Plus (Illumina), yielding 150-nucleotide paired-end reads.

### Virome assembly and virus identification

Raw sequencing reads were first assessed for quality using FastQC (Andrews 2010), followed by adapter removal and quality trimming with Trimmomatic (Bolger et al., 2014) to ensure high-quality input for assembly. Cleaned reads were then assembled *de novo* using MEGAHIT v1.2.9 (Li et al., 2015), which is optimised for complex and large metagenomic datasets and performs efficiently even on high-volume virome data (Li et al., 2015). Resulting contigs were annotated by searching protein-coding regions against NCBI nr database using DIAMOND blast v2.1.9 (Buchfink et al., 2015) using the non-redundant protein database obtained from the National Center for Biotechnology Information (NCBI), enabling rapid alignment and putative functional and taxonomic assignment of viral sequences (Hernández-Salmerón & Moreno-Hagelsieb, 2020).

To estimate the relative abundance of viruses across samples, quality-filtered reads were mapped back to assembled viral contigs using Bowtie2 v2.4.4 (Langmead & Salzberg, 2012), and abundance metrics were calculated from the mapping results, standardising to reads per million (RPM). Taxonomic information for viral contigs was derived from DIAMOND blast results and curated with available reference databases, enabling identification of viral families and potential novel viruses. Contig-level data, including contig abundance, taxonomy, and closest genetic matches, were compiled to support further analysis, including manual screening during which non-viral genetic matches were identified and removed (Chang et al., 2024). To minimise the risk of false positives, a E-value cut-off of 1 × 10^−10^ was applied during the BLAST searches.

### Virus phylogenetic analysis

To investigate the evolutionary relationships among the identified viruses and to infer their likely host associations, we constructed phylogenetic trees using conserved protein-coding genes. Viruses that clustered with known avian or vertebrate viruses were considered likely avian-associated, whereas those grouping with viruses associated with invertebrates, plants, fungi, bacteria or environmental sources were interpreted as originating from dietary or environmental sources, or sample processing contamination.

Translated viral protein sequences from conserved genes (e.g. RNA-dependent RNA polymerases (RdRp), DNA polymerases, replicase, non-structural proteins, or capsid proteins for non-polymerase encoding viruses) were aligned using MAFFT v7.490 (L-INS-i algorithm) with representative sequences from the same viral family available in NCBI RefSeq, along with the closest BLASTp genetic relatives (Katoh & Standley, 2013). Poorly aligned regions were removed using trimAl v1.5.0 (Capella-Gutiérrez et al., 2009) with a gap threshold of 0.9. Maximum likelihood phylogenies were estimated with IQ-TREE v2.4.0 (Nguyen et al., 2015), with the best-fit substitution model selected by the built-in ModelFinder (Kalyaanamoorthy et al., 2017) and branch support assessed with 1,000 ultrafast bootstrap replicates (Minh et al., 2013). Resulting trees were visualised and annotated in FigTree v1.4.4 (Rambaut, 2018) and edited in Adobe Illustrator. Only viruses inferred to directly infect avian hosts, based on their phylogenetic placement alongside other vertebrate viruses, were retained for further analyses while those clustering with invertebrate-or environment-associated lineages were excluded.

### Identification of non-viral microorganisms

Microbial communities (including bacteria, fungi, archaea and eukaryotic parasites) were identified in each library using CCmetagen (v1.4.3; Marcelino et al. 2020) with default parameters, which assigns taxonomic classifications to metatranscriptomic reads based on k-mer–assisted alignment against reference databases. Reads were quality-filtered prior to analysis to remove low-quality bases and sequencing artefacts, as described above (see Virome assembly and virus identification). Taxonomic assignments were summarised at the phyla level, and read counts were used to estimate relative abundance within each library. To focus on dominant microbial taxa, the ten most abundant microbial families were selected for each host species based on mean relative read abundance across libraries.

### Microbial and viral community abundance and diversity analysis

All statistical analyses and visualisations were performed in RStudio v2025.05.1 using the tidyverse v2.0.0 (Wickham et al., 2019) and ggplot2 v3.5.2 (Wickham, 2016) packages. The effects of host species, diet, social behaviour, endemic status and sampling year on community diversity and composition were assessed. Alpha diversity was quantified at the family level using richness (number of families detected) and the Shannon diversity index, which accounts for both richness and evenness. Differences among host categories were tested using Kruskal–Wallis tests (Kruskal & Wallis, 1952), followed by Dunn’s post-hoc tests (Dunn, 1964) with Bonferroni correction where appropriate.

Beta diversity was assessed using Bray-Curtis dissimilarity matrices calculated from family-level abundance data and visualised using non-metric multidimensional scaling (NMDS) (Clarke, 1993). Ordinations were generated using the phyloseq (v1.50.0) (McMurdie & Holmes, 2013) and scatterplot3d (v0.3-44) (Ligges & Maechler, 2003) packages. Differences in community composition were tested using Permutational Multivariate Analysis of Variance (PERMANOVA) implemented in the adonis2 function of vegan (v2.7-1) (Oksanen et al., 2025). Host species, diet, social behaviour, endemic status and sampling year were included simultaneously as explanatory variables in the PERMANOVA models, allowing the effect of each factor to be assessed while accounting for the other variables in the model. Pairwise comparisons were performed using pairwiseAdonis (v0.4.1) (Martinez Arbizu, 2020), with Bonferroni-adjusted p-values and a significance threshold of α = 0.05.

### Viral nomenclature

Viruses were considered putative novel species if they shared less than 90% amino acid identity within a conserved protein sequence, such as the RdRp for RNA viruses or other family-specific conserved proteins for DNA viruses (e.g. DNA polymerase). Provisional names were assigned to all novel viral sequences based on the host species in which they were detected and, where possible, the viral genus they were most closely related to. When a virus was detected in, or suspected of infecting multiple species, it was referred to by the general descriptor ‘passerine-virus’ rather than named after a specific host species.

## Results

### Sampled passerine species harbour extensive and largely undescribed viral diversity

Metatranscriptomic sequencing generated a mean of 82.2 million reads per library (median = 79.2 million), with all samples providing sufficient depth for virome analyses (Supplementary Figure S2). From 443 passerines sampled between 2021 and 2024, pooled into 52 libraries, we identified avian viruses across 14 viral families (Figure 2). Viruses were detected in all host species sampled. *Picornaviridae* and *Astroviridae* were the most widespread viral families, occurring in 44 and 39 libraries, respectively, followed by *Caliciviridae* (33 libraries) and *Hepeviridae* (29 libraries). Viral diversity varied substantially among host species, with silvereyes and house sparrows harbouring the greatest number of viral families (11 and 9, respectively) (Figure 2).

**Figure 2.**
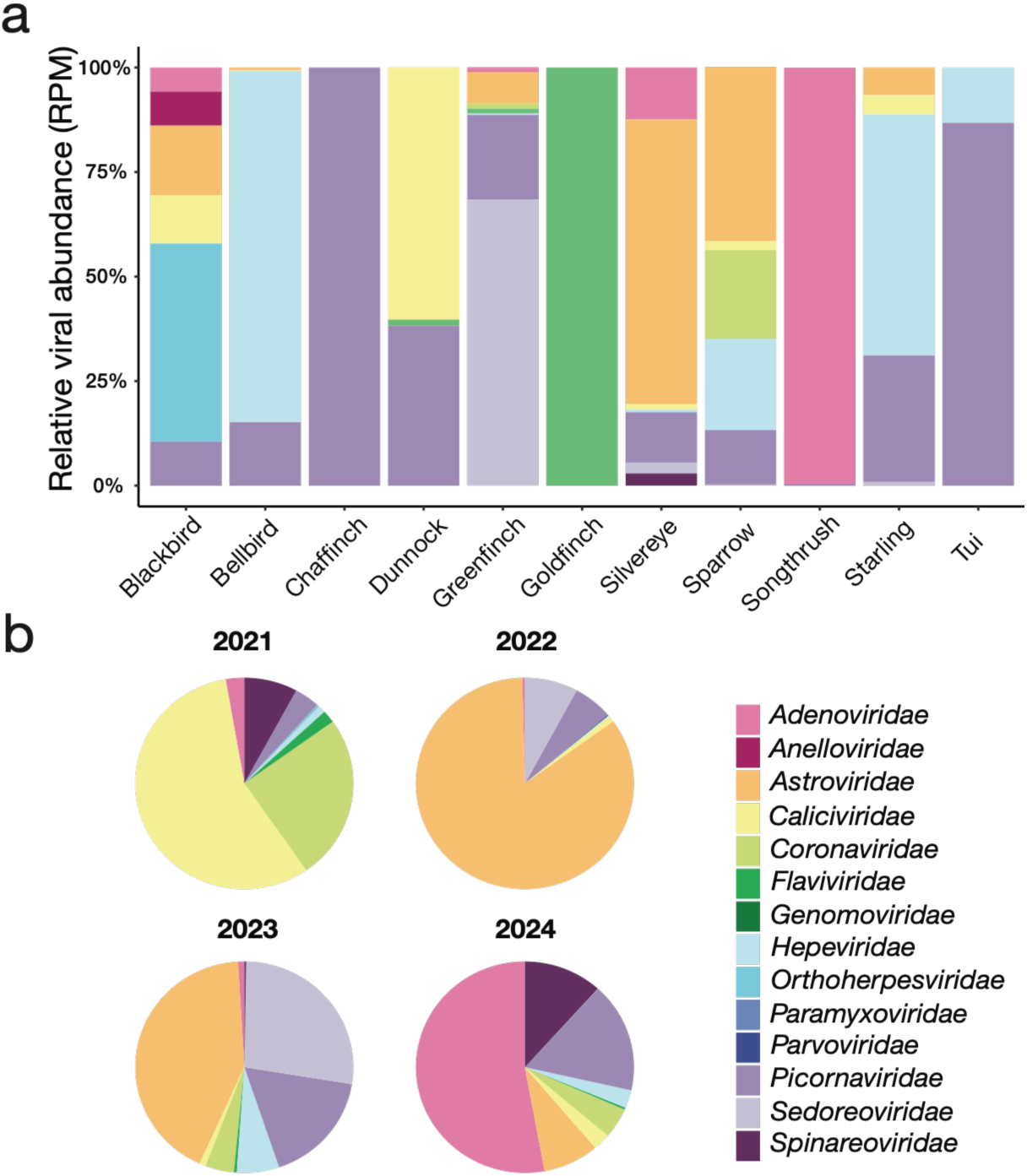
**(a)** Normalised relative abundance in reads per million (RPM) of each virus family detected across all host species sampled. **(b)** Pie charts illustrating the relative abundance of viruses found each sampling year (also in RPM). Colours in both panels correspond to the colour key shown.

### Evolutionary diversity of passerine viruses

Phylogenetic analyses identified 217 avian viruses, representing 69 unique species spanning 14 viral families (Supplementary Table S2). Of these 69 species, 54 (78%) were considered putatively novel, sharing <90% amino acid similarity with previously described viruses. These viruses encompassed diverse genome types, including 14 DNA viruses, three dsRNA viruses and 52 ssRNA viruses. Only avian-associated viruses were included in the analyses. All retained viruses clustered within established avian-associated lineages.

The *Picornaviridae* was the most diverse viral family detected, with 48 viruses representing 17 unique virus species distributed across multiple passerine hosts (Figure 3). These viruses spanned recognised genera, including Passerivirus and Hepatovirus, as well as several highly divergent lineages that could not be confidently assigned to existing genera. Many shared only 35-70% amino acid identity with their closest known relatives, including viruses previously identified incommon mynas (*Acridotheres tristis)* (Chang et al., 2021), robins (*Petroica australis)* (French, Stone, et al., 2022), grey-backed thrushes (*Turdus hortulorum)* (Woo et al., 2010), black swans (*Cygnus atratus*) (Shan et al., 2022) and other wild birds, indicating substantial undescribed diversity within this family.

**Figure 3.**
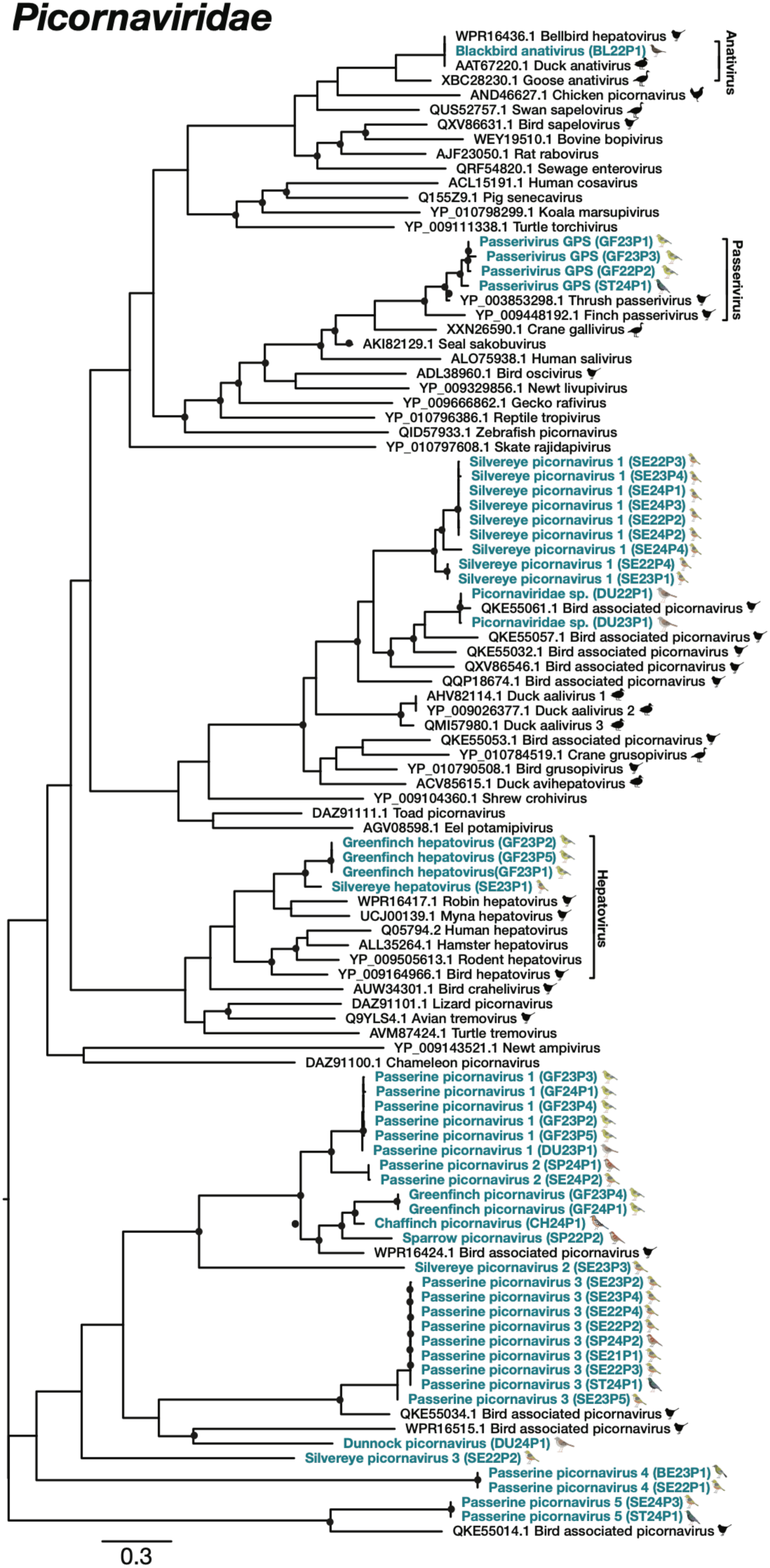
Maximum likelihood phylogenetic placement of RNA viruses detected in New Zealand passerines. The maximum likelihood tree was constructed using conserved RdRp genes from the *Picornaviridae* family. Passerine-derived viral sequences identified in this study are shown in teal and bold alongside representative reference sequences. Bird images beside the study sequences indicate the passerine host species in which each virus was detected, using the host symbols defined in Figure 1. Bird silhouettes beside reference sequences indicate previously described viruses of avian origin. The tree is midpoint-rooted for clarity, with branch lengths indicating the number of amino acid substitutions per site (scale bar). Nodes with bootstrap support >90% are marked with dots. This tree illustrates the evolutionary relationships between the passerine-associated picornaviruses identified in this study and previously characterised viruses.

The *Astroviridae* were similarly diverse, comprising 39 viruses representing 16 unique virus species within the genus *Avastrovirus* (Figure 4). Most novel astroviruses were closely related to viruses previously reported from Pallas’s rosefinches (*Carpodacus roseus*) (Shan et al., 2022) sharing approximately 82-89% amino acid identity, while two silvereye-associated astroviruses were more closely related to avian astroviruses. The *Caliciviridae* were also widespread, with eight virus species detected across several host taxa, most sharing only moderate amino acid similarity (44-70%) with previously described avian caliciviruses (Figure 4). Two deltacoronaviruses were detected in sparrows and greenfinches and shared 92-99% amino acid identity with Sparrow deltacoronavirus (Figure 4). Silvereyes harboured both avian paramyxovirus 8, which shared 95.2% amino acid identity with previously reported strains, and a novel, divergent paraavulavirus that shared only 57% amino acid identity with its closest known relative (Figure 4).

**Figure 4.**
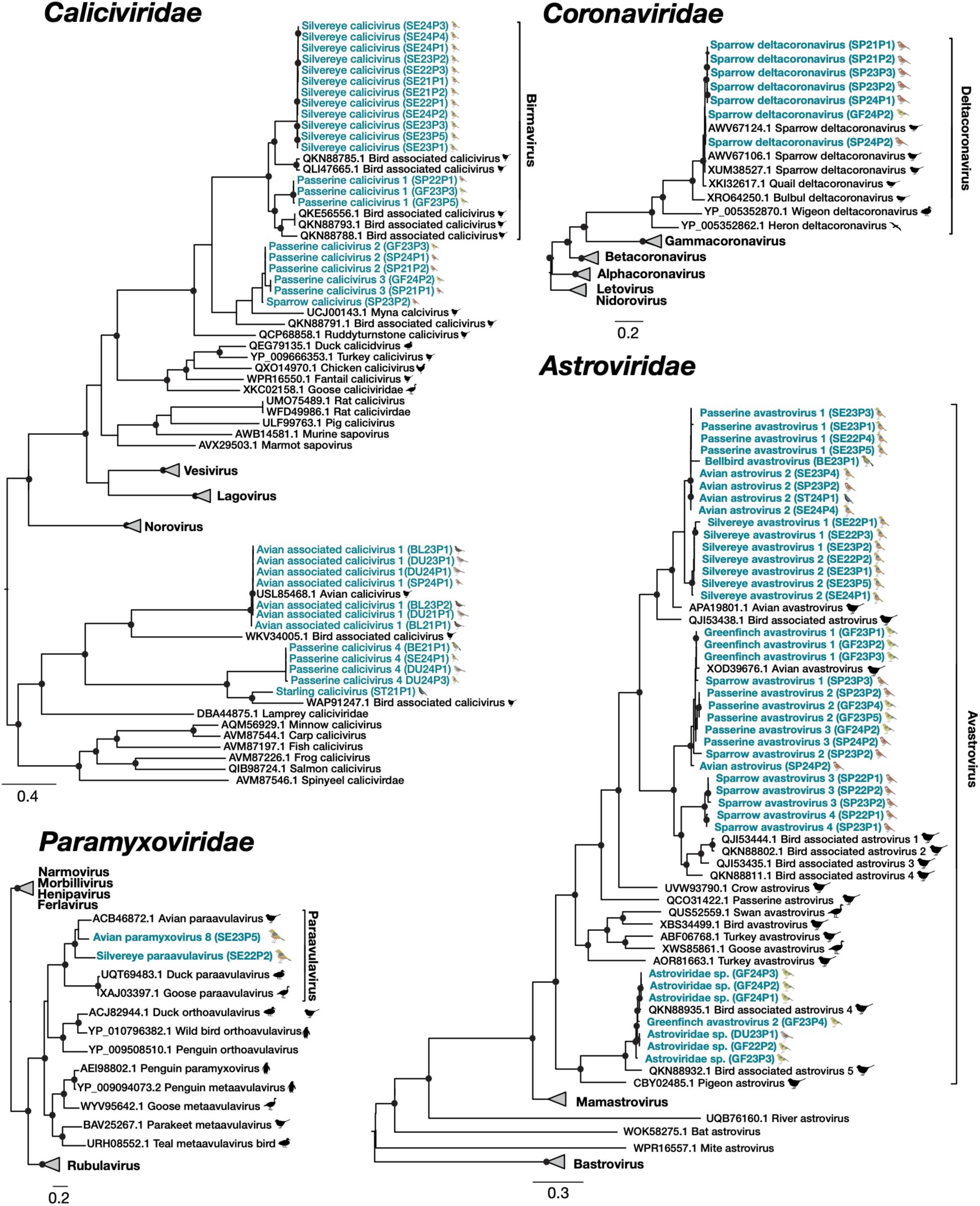
Maximum likelihood phylogenetic placement of RNA viruses detected in New Zealand passerines. Maximum likelihood trees were constructed using conserved RdRp genes from four virus families: *Caliciviridae, Coronaviridae, Paramyxoviridae,* and *Astroviridae.* Passerine-derived viral sequences identified in this study are shown in teal and bold alongside representative reference sequences from each family. Bird images beside the study sequences indicate the passerine host species in which each virus was detected, using the host symbols defined in Figure 1. Bird silhouettes beside reference sequences indicate previously described viruses of avian origin. Trees are midpoint-rooted for clarity, with branch lengths indicating the number of amino acid substitutions per site (scale bar). Nodes with bootstrap support >90% are marked with dots. These trees illustrate the evolutionary relationships between the passerine-associated viruses identified in this study and previously characterised viruses.

The dsRNA viruses comprised 15 rotaviruses representing two species (*Sedoreoviridae*) and 10 orthoreoviruses representing a single species (*Spinareoviridae*). Rotaviruses were detected in silvereyes, greenfinches and sparrows, including one lineage closely related to a rotavirus identified in velvet scoters (*Melanitta fusca*; 99% amino acid identity) and a novel rotavirus A lineage shared between greenfinches and sparrows that shared 85% identity with a rotavirus from rock pigeons (*Columba livia;* Figure 5). The orthoreovirus was detected in blackbirds and sparrows and clustered within established avian orthoreovirus diversity, sharing 99% amino acid identity with avian orthoreovirus (Figure 5).

**Figure 5.**
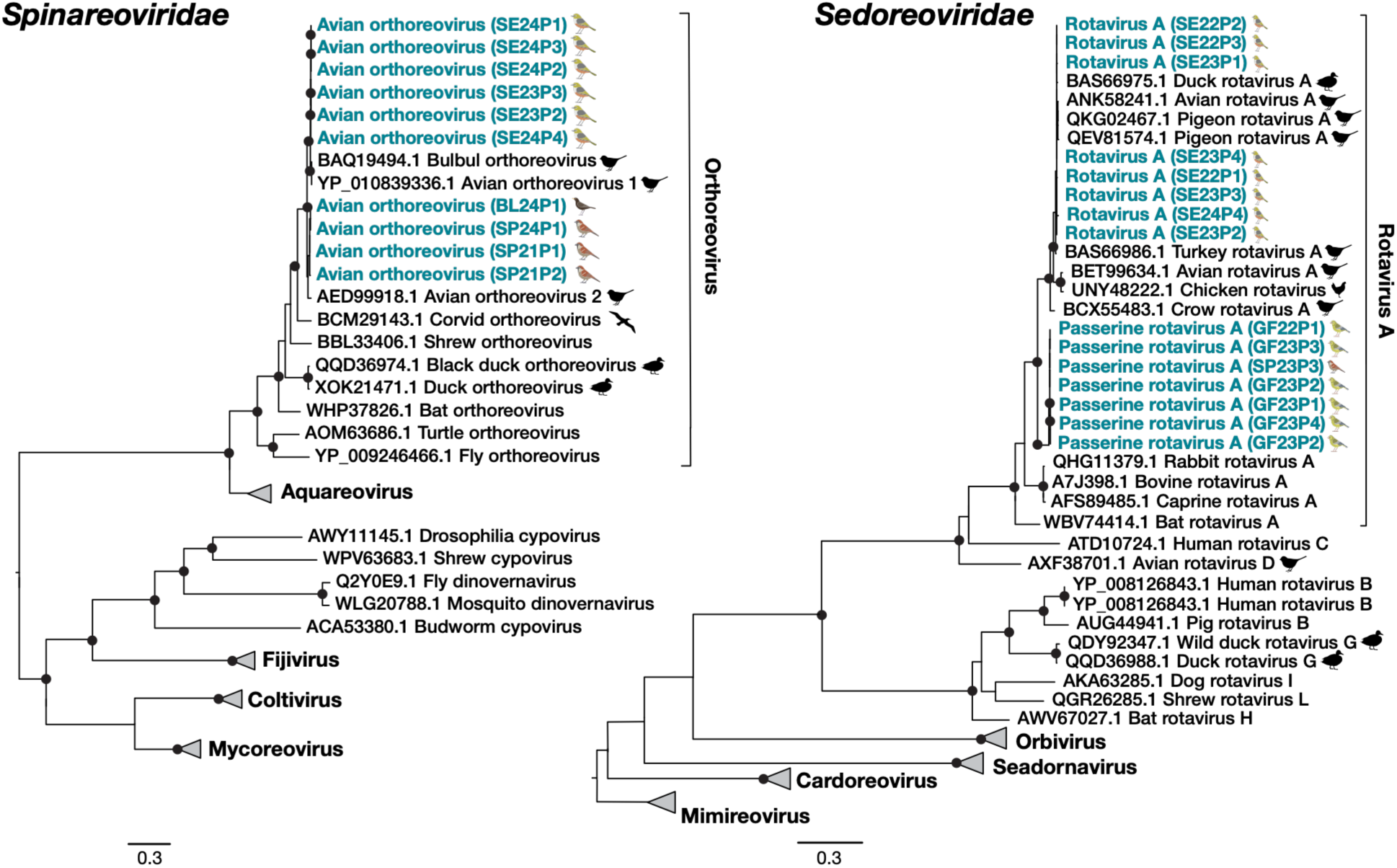
Maximum likelihood phylogenetic placement of dsRNA viruses detected in New Zealand passerines. Maximum likelihood trees were constructed using conserved RdRp genes from two dsRNA virus families: *Spinareoviridae* and *Sedoreoviridae*. Passerine-derived viral sequences identified in this study are shown in teal and bold alongside representative reference sequences from each family. Bird images beside the study sequences indicate the passerine host species in which each virus was detected, using the host symbols defined in Figure 1. Bird silhouettes beside reference sequences indicate previously described viruses of avian origin. Trees are midpoint-rooted for clarity, with branch lengths indicating the number of amino acid substitutions per site (scale bar). Nodes with bootstrap support >90% are marked with dots. These trees illustrate the evolutionary relationships between the passerine-associated viruses identified in this study and previously characterised viruses.

Hepeviruses and pegiviruses were also detected across multiple host species, including highly divergent lineages related to viruses previously identified in long-tailed tits (*Aegithalos caudatus)* (Lu et al., 2024), Eurasian tree sparrows (*Passer montanus*) and Brandt’s mountain finches (*Leucosticte brandti)* (Zhu et al., 2022), with amino acid identities ranging from 41–87% (Figure 6). Two hepeviruses were closely related to previously described avian viruses (>92% amino acid identity), whereas the remaining lineages were considerably more divergent, particularly Silvereye hepevirus, which shared only 41% amino acid identity with its closest known relative. Similarly, three of the four pegiviruses were considered novel, including a highly divergent passerine pegivirus detected in greenfinches that shared only 44% amino acid identity with *Passer montanus* pegivirus (Zhu et al., 2022) (Figure 6).

**Figure 6.**
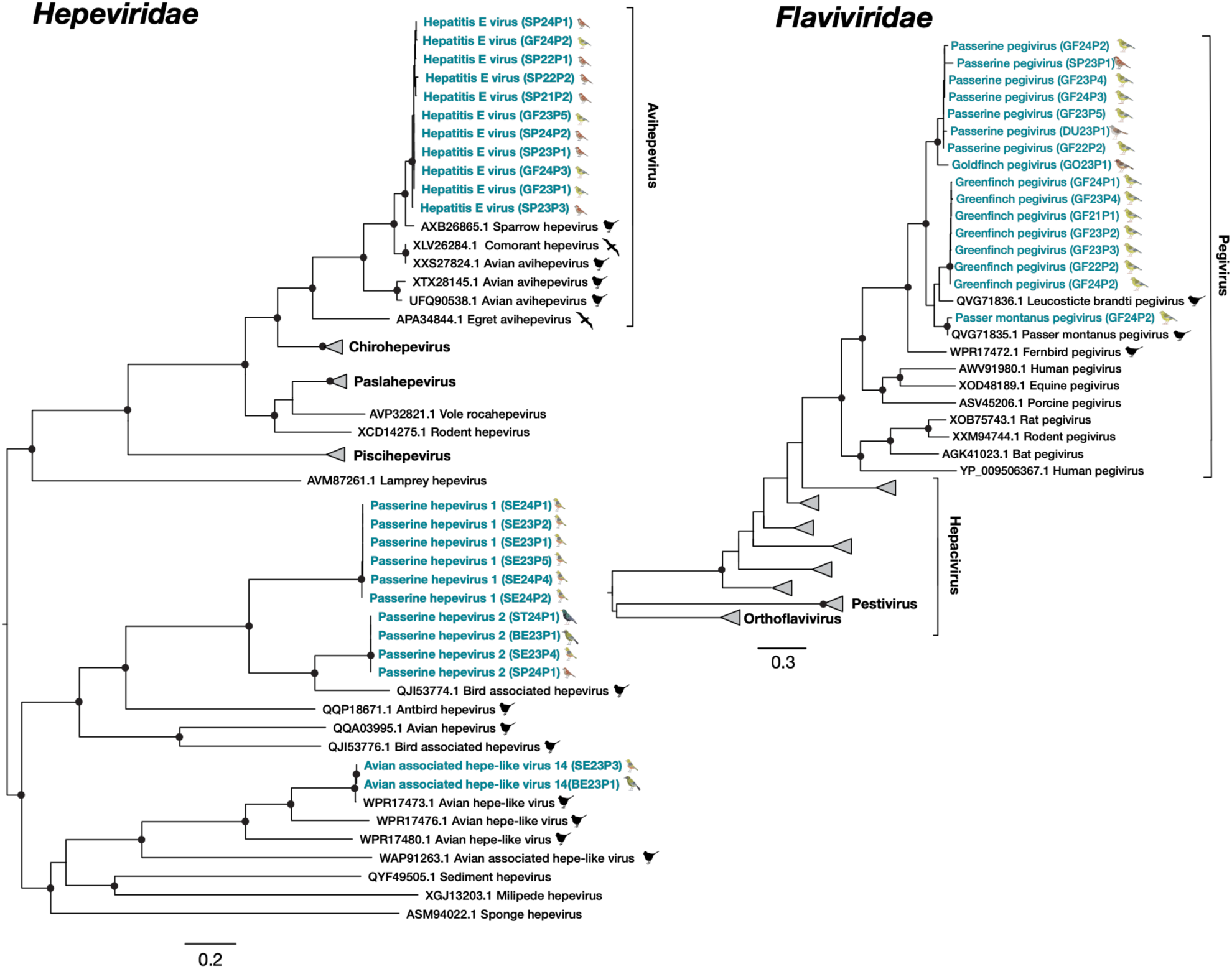
Maximum likelihood phylogenetic placement of RNA viruses detected in New Zealand passerines. Maximum likelihood trees were constructed using conserved RdRp genes from two virus families: *Hepeviridae* and *Flaviviridae*. Passerine-derived viral sequences identified in this study are shown in teal and bold alongside representative reference sequences from each family. Bird images beside the study sequences indicate the passerine host species in which each virus was detected, using the host symbols defined in Figure 1. Bird silhouettes beside reference sequences indicate previously described viruses of avian origin. Trees are midpoint-rooted for clarity, with branch lengths indicating the number of amino acid substitutions per site (scale bar). Nodes with bootstrap support >90% are marked with dots. These trees illustrate the evolutionary relationships between the passerine-associated viruses identified in this study and previously characterised viruses.

In contrast to the widespread distribution of many RNA viruses, several DNA viruses appeared host-restricted. Fourteen adenoviruses representing five unique species were identified, of which four species were putatively novel and clustered within established avian adenovirus lineages (Figure 7). These viruses were most closely related to adenoviruses previously reported from Naumann’s thrushes (*Turdus naumanni*) (Shan et al., 2022), grey warblers (*Gerygone igata*) (French et al., 2025) and domestic fowl (*Gallus gallus domesticus*) (Ojkic & Nagy, 2000), sharing 77-90% amino acid identity. Two novel alphaherpesviruses and three novel parvoviruses were detected exclusively in silvereyes and shared only 51-81% amino acid identity with their closest known relatives. Similarly, three novel gyroviruses were identified across blackbirds, silvereyes and sparrows, sharing 53-67% amino acid identity with previously described gyroviruses, while a novel bellbird genomovirus shared 71% identity with a finch-associated CRESS-DNA virus (Figure 7).

**Figure 7.**
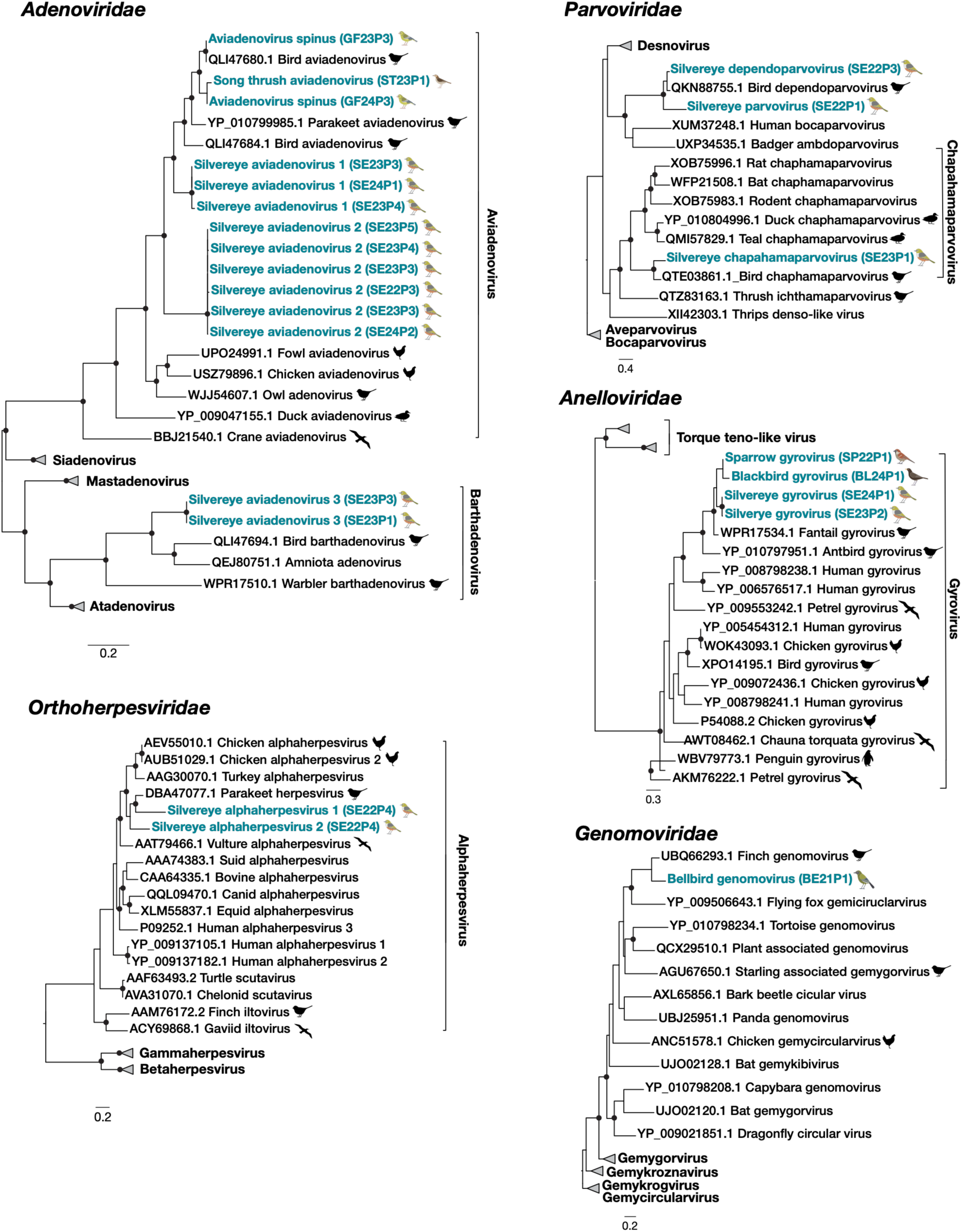
Maximum likelihood phylogenetic placement of DNA viruses detected in New Zealand passerines. Phylogenetic trees were constructed from conserved, family-specific genes: *Adenoviridae* (DNA polymerase), *Orthoherpesviridae* (DNA polymerase), *Parvoviridae* (NS1), *Anelloviridae* (ORF1/VP1), and *Genomoviridae* (capsid protein). Passerine-derived viral sequences identified in this study are shown in teal and bold alongside representative reference sequences from each family. Bird images beside the study sequences indicate the passerine host species in which each virus was detected, using the host symbols defined in Figure 1. Bird silhouettes beside reference sequences indicate previously described viruses of avian origin. Trees are midpoint-rooted for clarity. Branch lengths represent the number of amino acid substitutions per site (scale bar). Nodes with bootstrap support greater than 90% are indicated with black circles. These trees illustrate the evolutionary relationships of the viruses detected in this study relative to previously characterised viruses.

Across all viral families, amino acid similarity to the closest known relatives ranged from approximately 35% to 99%, highlighting the broad evolutionary diversity of passerine-associated viruses in New Zealand. Many viruses were most closely related to viruses previously reported from wild birds in Asia, Australia, Europe and North America, whereas others formed deeply divergent lineages with no close relatives currently available in public databases, indicating that a substantial proportion of avian viral diversity remains unsampled.

### Host species and sociality structure passerine viromes

To determine the ecological factors associated with variation in passerine viromes, we compared viral richness, diversity and community composition among host species and across ecological traits, including social behaviour, diet, endemic status and sampling year. We found that host species was the strongest determinant of virome composition, explaining 33.1% of the variation in community structure (PERMANOVA: R² = 0.33, *p* = 0.001) (Figure 8). Viral richness also differed significantly among species, with the greatest differences observed between silvereyes and dunnocks (adjusted *p* = 0.0054) (Supplementary Figure S3).

**Figure 8.**
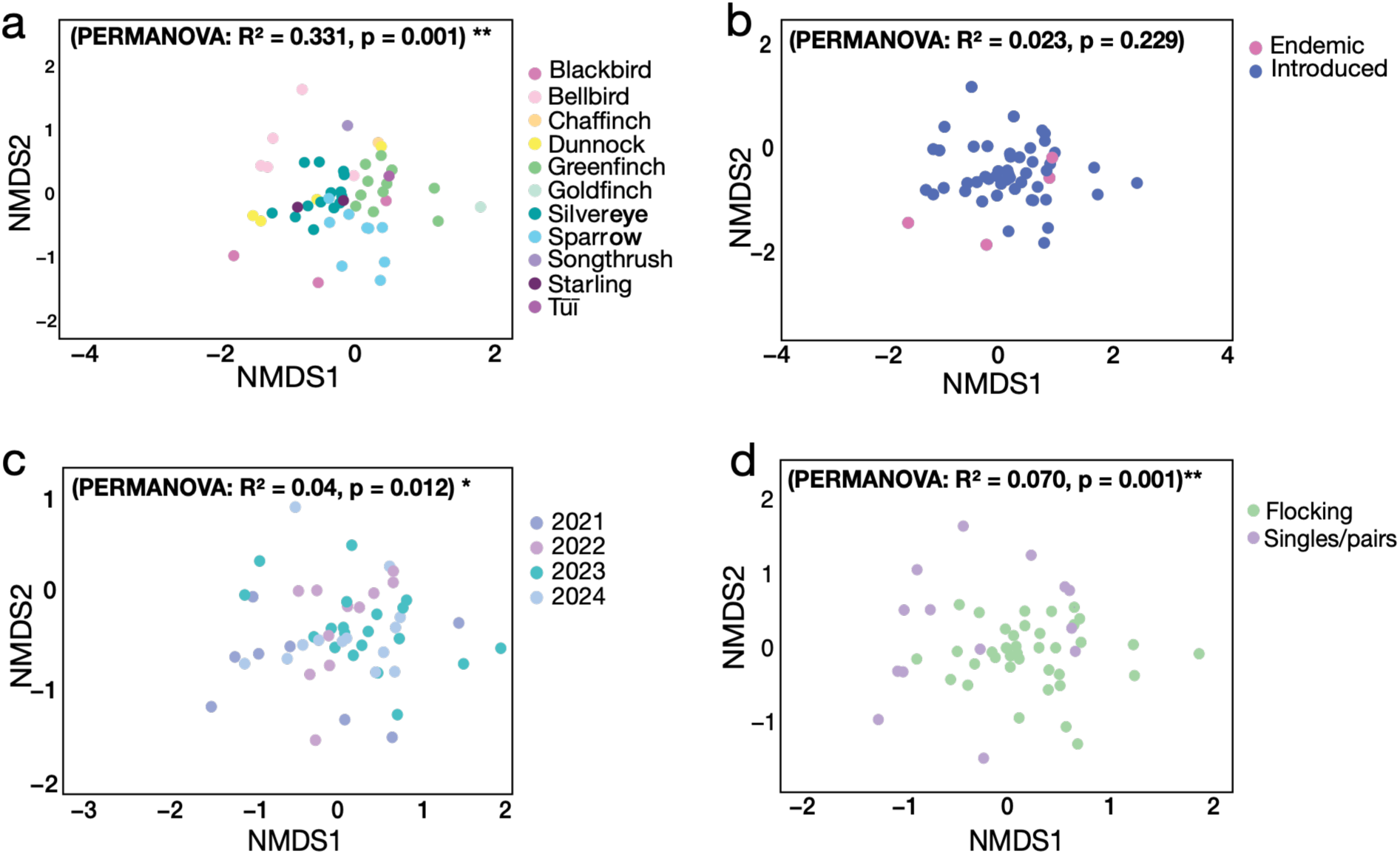
NMDS ordinations of passerine viral community composition. Non-metric multidimensional scaling (NMDS) ordinations depict viral community composition based on Bray–Curtis dissimilarities. Stress = 0.195. Panels illustrate differences in viral family composition across **(a)** host species, (**b)** introduction status (endemic vs introduced), **(c)** sampling year, and **(d)** social category.

We next tested whether social behaviour influenced viral diversity, as flocking species experience more frequent and prolonged contact among conspecifics than solitary or pair-living species, potentially increasing opportunities for viral transmission within populations. Flocking species harboured significantly greater viral richness (Kruskal–Wallis: p < 0.00001) and Shannon diversity (*p* = 0.03) than solitary or pair-living species. Social behaviour explained 7% of the variation in virome composition (PERMANOVA: R² = 0.070, p = 0.001), making it the strongest ecological predictor after host species (Figure 8).

To assess whether host endemic status influenced virome structure, we compared endemic and introduced species. Introduced species exhibited higher viral richness and diversity than endemic species (richness: adjusted *p* = 0.02; Shannon diversity: *p* = 0.02) (Supplementary Figure S3), although endemic status explained little variation in overall community composition (PERMANOVA: R² = 0.023, p = 0.23). Finally, we tested whether dietary niche influenced virome composition, as species consuming different food resources may encounter distinct viral reservoirs and transmission pathways. Birds were classified into their broad groups based on their predominant diet (insectivores, granivores, omnivores and nectarivores), but diet was not associated with virome composition. In contrast, sampling year explained approximately 4% of the variation in community structure (PERMANOVA: R² = 0.040, p = 0.012), suggesting modest temporal variation in viral communities between years. However, differences in sampling effort and host species representation among years, particularly for silvereyes and greenfinches (Figure 1), may have contributed to this variation.

### Broad overlap in viral family detection among host species

Host-virus family network analyses revealed broad overlap in the viral families detected among passerine species (Figure 9). *Picornaviridae* and *Astroviridae* were widely distributed across hosts and accounted for the majority of host–viral-family associations, while *Caliciviridae* and *Hepeviridae* were also detected in multiple host species. In contrast, several DNA virus families, including *Genomoviridae* and *Orthoherpesviridae*, were restricted to a single host species. Network connectance was moderate (35.7%), meaning that just over one-third of all possible host–viral-family associations were observed, while linkage density was 2.40. Nestedness was low (Nestedness metric based on Overlap and Decreasing Fill; NODF = 11.1 on a 0–100 scale), indicating that the viral families detected in hosts with fewer associations were not consistently subsets of those detected in more highly connected hosts. Shannon diversity of interactions was 2.10, with an evenness of 0.52, indicating that associations were unevenly distributed among host species and viral families. Host species were associated with a mean of five viral families, while each viral family was detected in a mean of 3.93 host species. The silvereye was the most highly connected host, with associations to 11 viral families. Community detection identified four modules, although modularity was low (Q = 0.21), indicating weak clustering and substantial overlap in viral-family assemblages among hosts.

**Figure 9.**
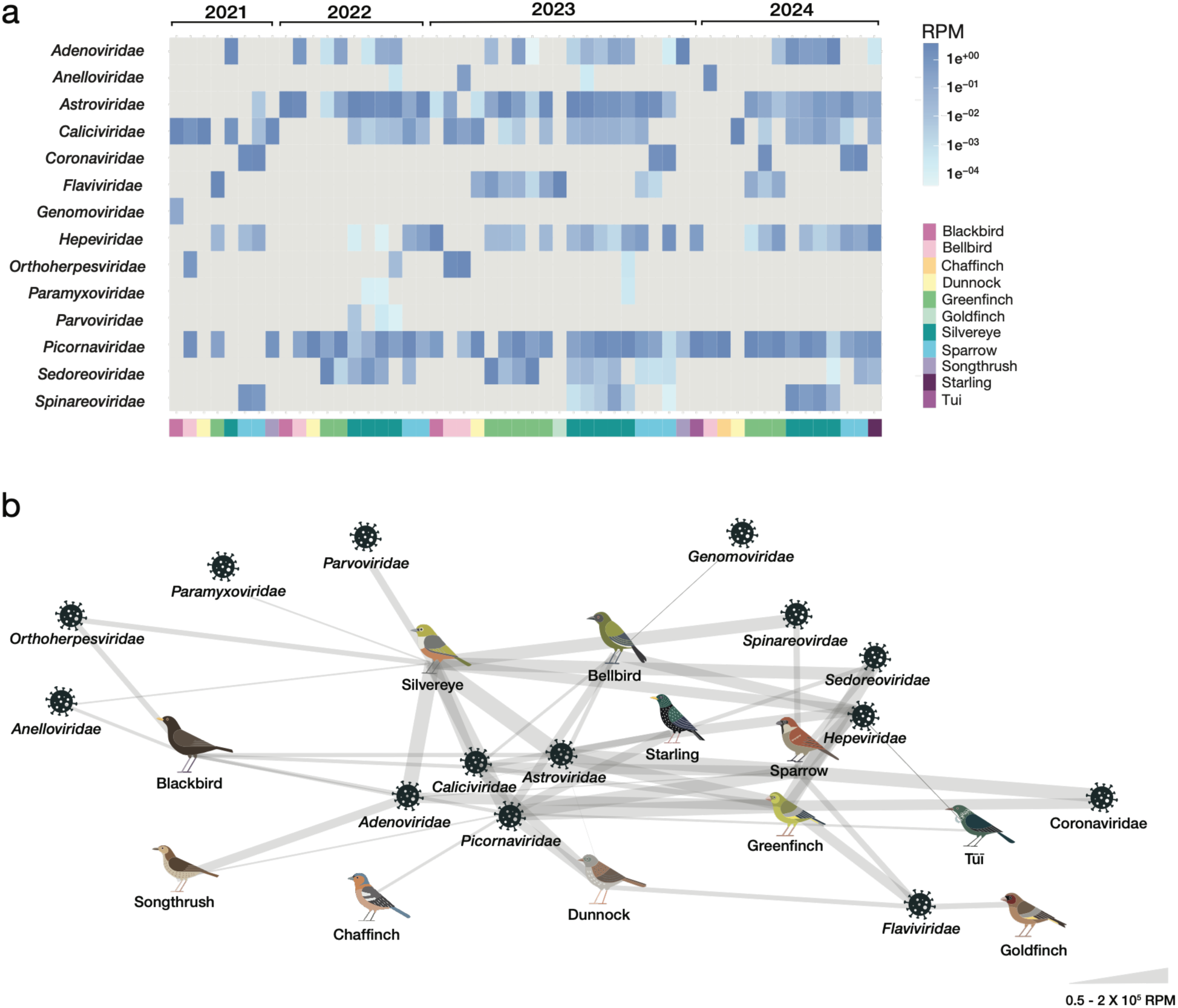
Passerine virome composition and host–virus interaction networks. **(a)** Heatmap of normalised viral abundances (reads per million, RPM) for viral families across passerine libraries. The colour gradient from white to blue represents increasing viral abundance, with white indicating absence of sequencing reads and darker blue indicating high relative abundance. Columns correspond to sequencing libraries, with collection year indicated above, and rows represent viral families. Host species are shown as coloured annotation blocks below the heatmap. (b) Network analysis of viral families detected across passerine bird species. Each node represents either a bird species (illustrated by bird icons) or a viral family (grey virus-shaped nodes). Edges connect host species to viral families, with edge thickness proportional to the relative abundance of sequencing reads mapping to that association (in reads per million, RPM). A scale for edge thickness is provided.

### Viral and microbial communities respond to different ecological drivers

To determine whether the ecological factors shaping passerine viromes also influenced non-viral host-associated microbial communities (bacteria, fungi, protists, etc.), we performed equivalent community diversity analyses on the non-viral microbiome. Microbial community composition differed significantly among host species (PERMANOVA: R² = 0.45, *p* = 0.006) and was also influenced by diet (R² = 0.16, *p* = 0.03) (Figure 10). In contrast, social behaviour, endemic status and sampling year were not significant predictors of non-viral microbial community structure. The factors shaping microbial and viral communities therefore differed substantially. Whereas social behaviour was a strong predictor of viral richness, diversity and community composition, non-viral microbial communities were structured primarily by host identity and diet, although substantial variation remained unexplained.

**Figure 10.**
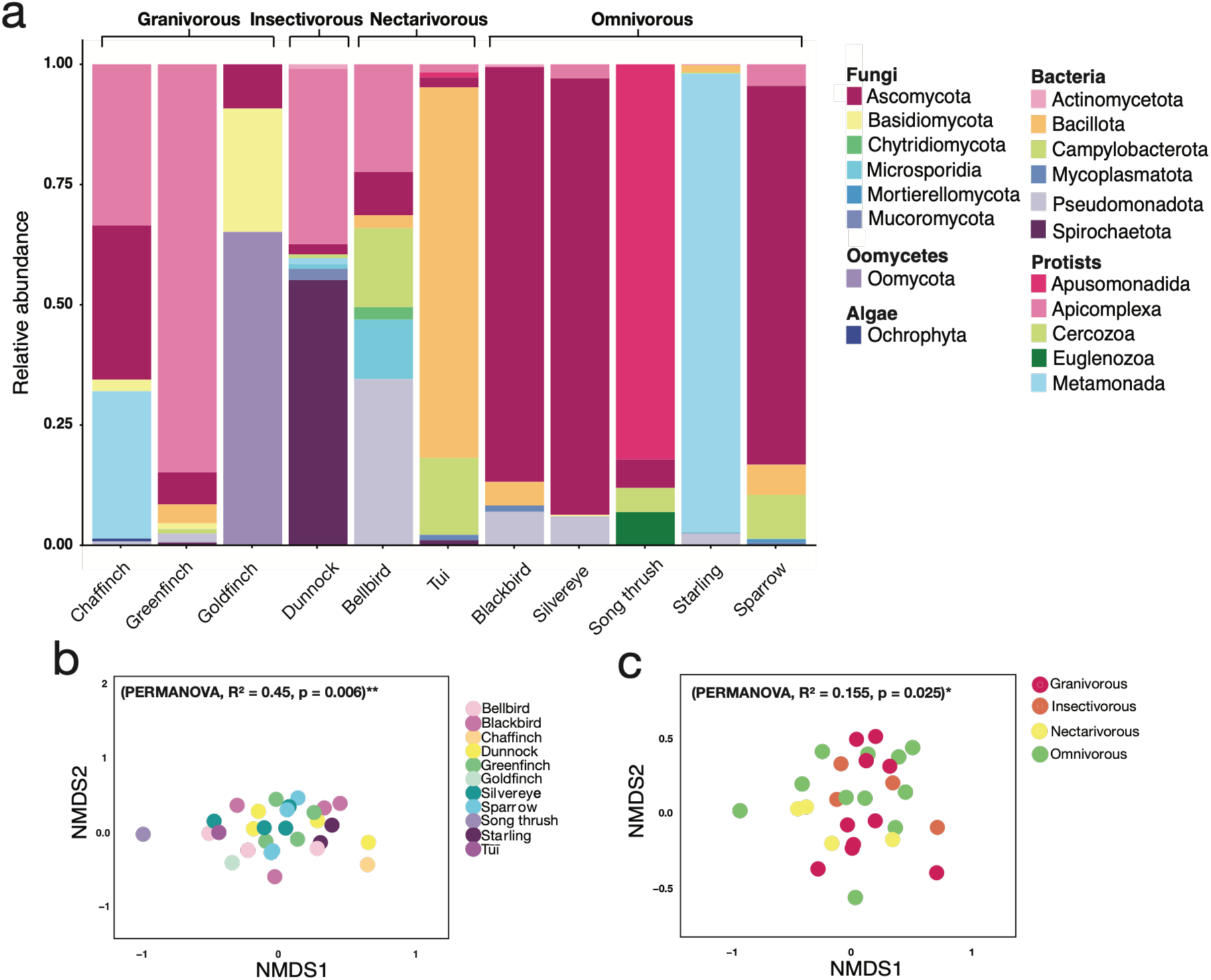
Microbial community composition by host species and diet. **(a)** Stacked bar plot showing relative abundance of the ten most abundant microbial phyla across bird species. **(b-c)** Non-metric multidimensional scaling (NMDS) ordinations of microbial communities coloured by host species **(b)** and predominant dietary group **(c)**, based on Bray–Curtis dissimilarity. Stress = 0.228.

## Discussion

We have characterised the viromes of 443 passerines from Aotearoa New Zealand and identified 69 avian viruses, including 54 putatively novel species. These findings demonstrate that New Zealand passerines harbour substantial previously undescribed viral diversity. The presence of numerous viruses in apparently healthy individuals further reinforces the capacity of wild birds to harbour diverse viral communities and highlights the extent to which avian viral diversity remains under-sampled globally (Chang et al., 2020; Wille et al., 2018).

Host species was the strongest determinant of virome composition, explaining approximately one-third of the observed variation. This strong association between host and virome composition is consistent with previous studies across multiple host classes, which have identified marked differences in virome composition among taxonomically divergent species, including in passerines, despite species occurring within the same ecosystem (Chen et al., 2023; Geoghegan et al., 2021; Grimwood et al., 2024). Similarly, studies of wild waterfowl and shorebirds have identified substantial virome heterogeneity among species within multi-species aggregations (Wille et al., 2019). These studies indicate that host species identity is an important determinant of virome composition even when opportunities for environmental exposure and interspecific contact overlap.

Among the ecological factors examined, social behaviour emerged as the strongest predictor of viral diversity. Flocking species harboured significantly greater viral richness and diversity than solitary or pair-living species, consistent with theoretical expectations that increased contact rates facilitate pathogen transmission and persistence (Root et al., 2022). Social species form dense and highly connected transmission networks that increase opportunities for both within- and between-species spread, allowing a broader range of viruses to be maintained within populations (Webber et al., 2017; Willoughby et al., 2017). Social behaviour has been identified as an important factor influencing viral transmission and sharing in other vertebrate systems. For example, bats that co-roosted in the same caves were more likely to share viruses, providing evidence that opportunities for close association between host species can influence patterns of viral sharing (Willoughby et al., 2017). Our results suggest that similar processes may structure viromes in wild passerines, with greater social connectivity potentially increasing opportunities for viral acquisition and maintenance.

In the present study, evidence of viral sharing among host species was widespread. Several RNA virus families, including astroviruses, caliciviruses, picornaviruses and hepeviruses, were detected across multiple passerine species, consistent with broader host associations and greater potential for cross-species transmission. In contrast, several DNA virus families were restricted to individual host species, consistent with greater host specificity, although the evolutionary processes underlying these associations cannot be resolved from the present data. These contrasting patterns support the view that RNA viruses generally occupy broader host ranges and transmit more readily among hosts than DNA viruses (Geoghegan et al., 2017; Reuther et al., 2021; Tian et al., 2022). Notably, silvereyes occupied a central position within the virus-sharing network, harbouring numerous viral families also detected in other passerines. Their broad habitat use and gregarious behaviour suggest they may function as important hosts facilitating viral connectivity within passerine communities (Caron et al., 2015).

Many of the viruses detected here were closely related to viruses previously reported from Australia, Asia, Europe and North America (Dai et al., 2024; Lu et al., 2024; Ojkic & Nagy, 2000; Shan et al., 2022). Despite New Zealand’s geographic isolation, its avifauna has strong evolutionary links with the wider Australasian region, including several passerine lineages with Australasian origins (Gibb et al., 2015). The presence of closely related viruses across geographically distant regions may therefore reflect a combination of shared host evolutionary history and viral dispersal, although the relative contribution of these processes cannot be determined from the present data. Conversely, the detection of highly divergent viruses suggests that local viral diversification has also contributed to the observed virome.

Comparison with broader microbial communities revealed that viruses and other microbes may be influenced by different ecological processes. Whereas virome composition was strongly influenced by social behaviour, microbial community structure was more strongly associated with host identity and diet. However, the measured factors explained only a relatively small proportion of the variation in microbial community structure, indicating that much of this variation may reflect unmeasured environmental, temporal or individual-level factors. This comparison should also be interpreted cautiously because the virome analysis was restricted to viruses classified as avian, whereas the microbial analysis included the broader microbial community. These dietary categories represent broad feeding strategies, as passerine diets can vary seasonally and often span multiple food types. Similar differences in the ecological factors shaping viral and broader microbial communities have been reported in other host systems, suggesting that viruses may be influenced by ecological processes distinct from those structuring other components of the microbiome (Graciette et al., 2023; Yao et al., 2024). Differences in detection sensitivity between viral and microbial communities may further limit direct comparisons. Nevertheless, the association between viral community composition and social behaviour is consistent with the role of host contact in viral transmission, where direct interactions and shared resources or environments can provide opportunities for viral exposure and transmission (Root et al., 2022).

These findings should be interpreted in the context of the opportunistic sampling design, which resulted in unequal numbers of birds sampled across host species and years. Sampling was also conducted at a single backyard site, limiting the extent to which the findings represent passerine viromes across other locations or habitats. As individuals were pooled by species and year before sequencing, some species–year combinations were represented by only one library, limiting replication and making the effects of host species and sampling year difficult to disentangle. The yearly patterns shown in Figure 2b may therefore partly reflect which species were sampled in each year, rather than genuine temporal variation in viral diversity. Pooling enabled the cost-effective screening of 443 birds but prevented viruses from being associated with individual birds and may have reduced the detection of uncommon or low-abundance viruses through dilution. These aspects of the sampling design may have contributed to the observed patterns of viral richness and diversity.

In this study, we show that passerine viromes are structured by a combination of host identity and social behaviour, with flocking species supporting richer and more diverse viral communities than less social hosts. The extensive sharing of viruses among species further suggests that ecological interactions such as through shared food and water sources may create pathways for viral transmission across host boundaries. These findings have broader implications for understanding how viruses circulate within complex avian communities, particularly where introduced and endemic species coexist and share habitats and resources. This is especially relevant in New Zealand, where the susceptibility of many native bird species to emerging viruses remains poorly understood. The recent detection of H5N1 highly pathogenic avian influenza virus in New Zealand provides a timely example. Although long-distance HPAI dispersal is primarily associated with migratory waterbirds, passerines can also become infected and may contribute to local transmission or cross-species viral exchange (Fujimoto et al., 2015). Our finding that social behaviour is strongly associated with viral diversity highlights the potential importance of highly connected and gregarious species in facilitating viral circulation within bird communities. More broadly, understanding which host traits promote viral diversity and sharing can help identify species and ecological interfaces most likely to facilitate the establishment and spread of newly introduced viruses, informing surveillance and conservation responses to future disease threats.

## Data availability

Sequencing reads are available on NCBI’s sequence read archive (SRA) under BioProject PRJNA1484759 while all virus sequences are available on GenBank under accessions PZ720416-PZ720632.

## Author contributions

Jemma Geoghegan, Benjamin Perry and Bruce Robertson designed the study. Kristina Hames, Janelle Wierenga, Benjamin Perry, Bruce Robertson and Fiona Robertson conducted the fieldwork and collected samples. Kristina Hames performed the laboratory work, led the bioinformatic analyses and prepared the first draft of the manuscript. Stephanie Waller and Rebecca French supervised the laboratory and bioinformatic analyses. Jemma Geoghegan secured funding and provided overall supervision. All authors contributed to reviewing and editing the manuscript and approved the final version.

