## Supplementary figures and images for "Social behaviour is a key determinant of virome diversity in New Zealand passerines"

### Supplementary Figure 1

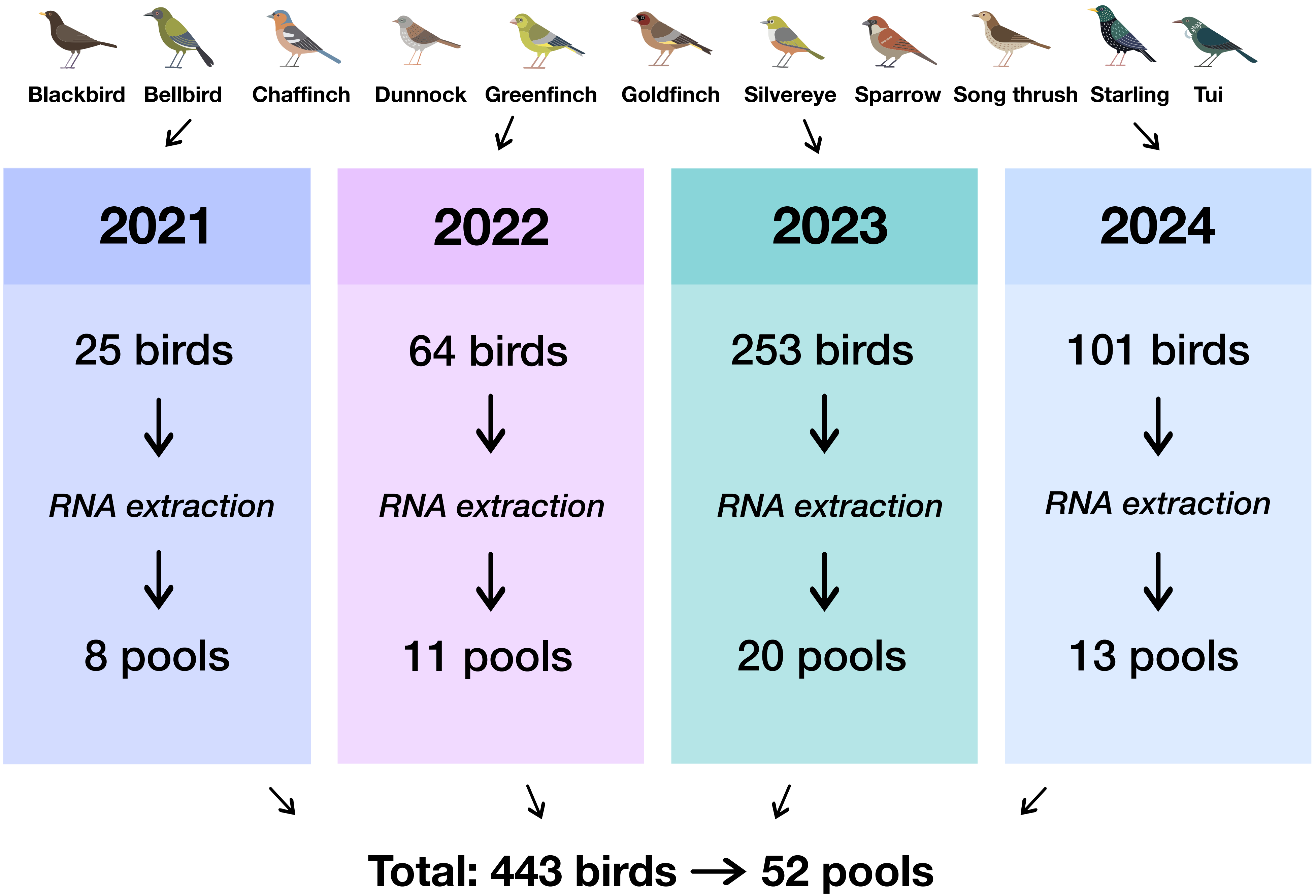

### Supplementary Figure 2

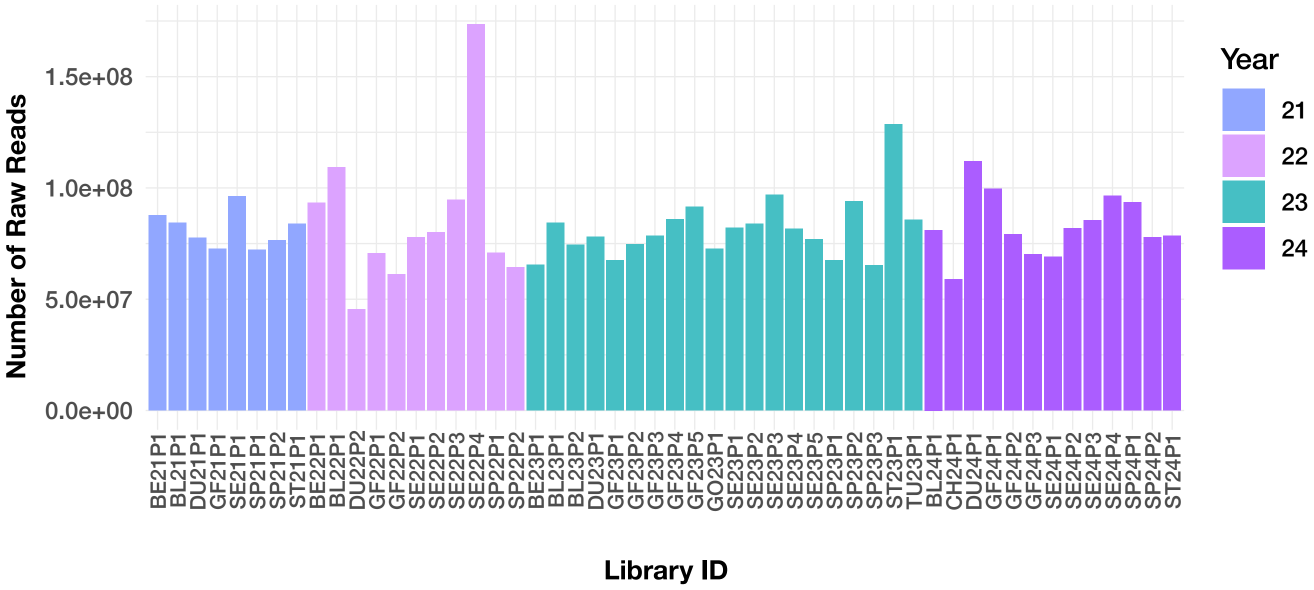

### Supplementary Figure 3

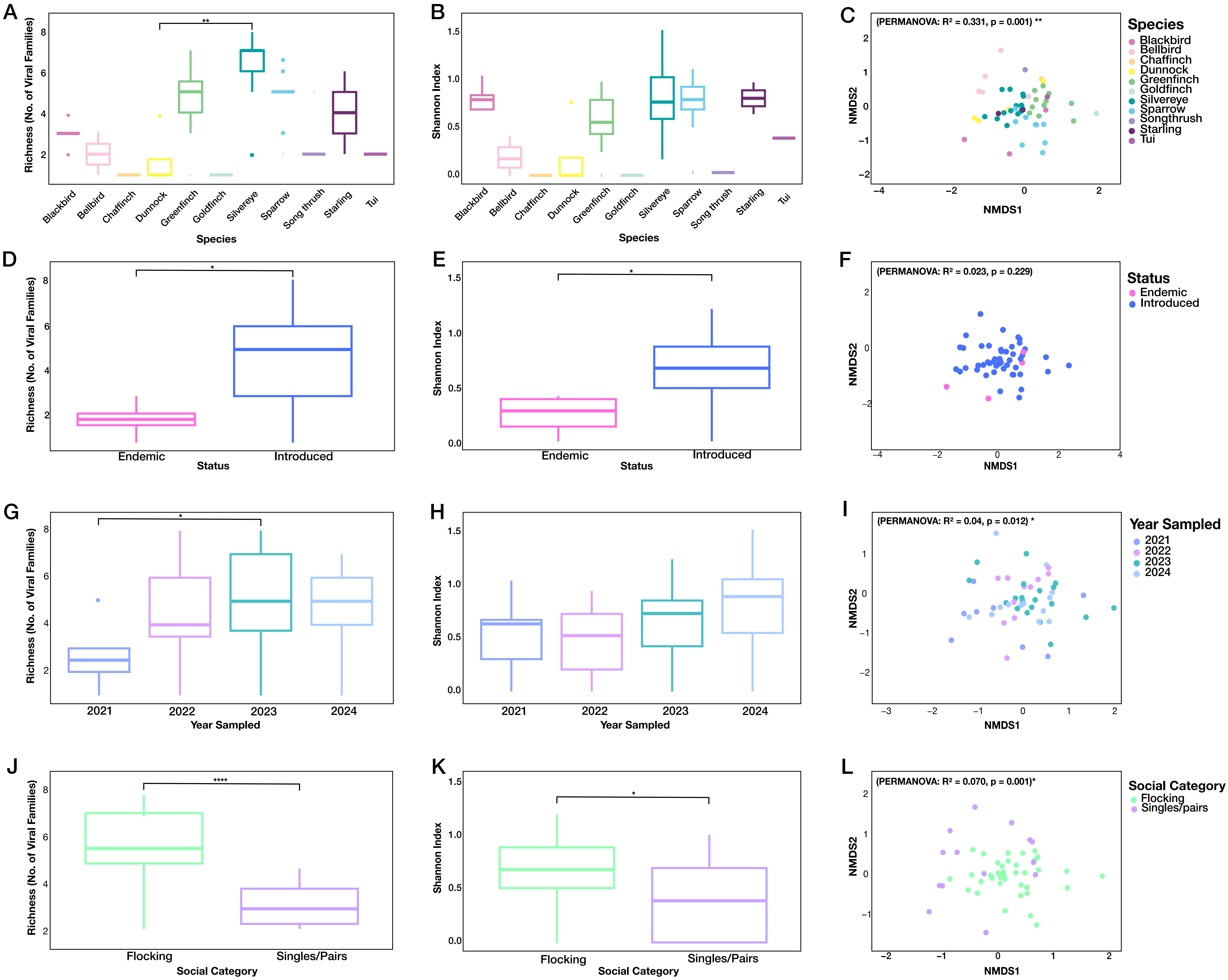
