## Supplementary Table 1 for "Social behaviour is a key determinant of virome diversity in New Zealand passerines"

**Supplementary Table S1.** Passerine bird species sampled between 2021 and 2024. The number of individuals of each species caught per year, along with their origin, predominant diet, and social behaviour. Origins are defined as: endemic = species that evolved in New Zealand; self-introduced = species that colonised naturally and established populations; introduced = species brought to New Zealand by humans. Predominant diet categories are simplified as insectivorous, granivorous, nectarivorous, or omnivorous, recognising that most species show dietary flexibility. Predominant social behaviour is classified as: flocking = species that commonly form groups outside the breeding season; singles/pairs = species that primarily occur alone or in pairs.

| **Species** | **Species Origin** | **Predominant Diet** | | **Social Behaviour** | **2021** | **2022** | **2023** | **2024** | **Total** |
| --- | --- | --- | --- | --- | --- | --- | --- | --- | --- |
| **Bellbird**  (*Anthornis melanura*) | Endemic | Omnivorous | | Single/pairs | 2 | 1 | 4 | 0 | 7 |
| **Blackbird**  *(Turdus merula*) | Introduced | Omnivorous | | Single/pairs | 2 | 2 | 8 | 2 | 14 |
| **Chaffinch**  (*Fringilla coelebs*) | Introduced | Granivorous | | Flocking | 0 | 0 | 0 | 1 | 1 |
| **Dunnock**  (*Prunella modularis*) | Introduced | Insectivorous | | Single/pairs | 2 | 1 | 7 | 1 | 11 |
| **Goldfinch**  (*Carduelis carduelis*) | Introduced | Granivorous | | Flocking | 0 | 0 | 1 | 0 | 1 |
| **Greenfinch**  (*Chloris chloris*) | Introduced | Granivorous | | Flocking | 4 | 9 | 103 | 38 | 154 |
| **House Sparrow**  (*Passer domesticus*) | Introduced | Omnivorous | | Flocking | 10 | 8 | 20 | 12 | 50 |
| **Silvereye**  (*Zosterops lateralis*) | Self-introduced | Omnivorous | | Flocking | 4 | 43 | 102 | 44 | 193 |
| **Song Thrush**  (*Turdus philomelos*) | Introduced | Omnivorous | | Single/pairs | 0 | 0 | 1 | 0 | 1 |
| **Starling**  (*Sturnus vulgaris*) | Introduced | Omnivorous | | Flocking | 1 | 0 | 0 | 3 | 4 |
| **Tūī** (*Prosthemadera* *novaeseelandiae*) | Endemic | Nectarivorous | | Single/pairs | 0 | 0 | 7 | 0 | 7 |
| **Total** |  | |  |  | 25 | 64 | 253 | 101 | 443 |
