## Supplementary Table 2 for "Social behaviour is a key determinant of virome diversity in New Zealand passerines"

Supplementary Table S2. Detailed information on viruses detected in this study.

Metadata include the virus name, corresponding host species, and the closest known genetic relatives based on sequence similarity.

| Virus Name | Family | Host | Top Hit Name | Amino Acid Identity |
| --- | --- | --- | --- | --- |
| Aviadenovirus spinus | *Adenoviridae* | Greenfinch | *Aviadenovirus spinus* | 99.09% |
| Silvereye adenovirus | *Adenoviridae* | Silvereye | *Adenoviridae* sp. | 71.63% |
| Silvereye aviadenovirus 1 | *Adenoviridae* | Silvereye | *Adenoviridae* sp. | 80.12% |
| Silvereye aviadenovirus 2 | *Adenoviridae* | Silvereye | *Fowl aviadenovirus* 4 | 77.06% |
| Song thrush aviadenovirus | *Adenoviridae* | Song thrush | *Adenoviridae* sp. | 89.47% |
| Blackbird gyrovirus | *Anelloviridae* | Blackbird | *Fantail gyrovirus* | 66.67% |
| Silvereye gyrovirus | *Anelloviridae* | Silvereye | [*Gyrovirus 10*](https://www.ncbi.nlm.nih.gov/Taxonomy/Browser/wwwtax.cgi?id=2218660) | 57.45% |
| Sparrow gyrovirus | *Anelloviridae* | Sparrow | *Gyrovirus 11* | 52.63% |
| Astroviridae sp. | *Astroviridae* | Greenfinch,Dunnock | *Astroviridae sp.* | 98.8% |
| Avian astrovirus | *Astroviridae* | Silvereye | *Avian astrovirus* | 90.00% |
| Avian astrovirus 1 | *Astroviridae* | Sparrow | *Avian astrovirus 1* | 90.68% |
| Avian astrovirus 2 | *Astroviridae* | Silvereye, Sparrow, Starling | *Avian astrovirus 2* | 99.52% |
| Bellbird avastrovirus | *Astroviridae* | Bellbird | *Astroviridae* sp. | 84.38% |
| Greenfinch avastrovirus 1 | *Astroviridae* | Greenfinch | *Astroviridae* sp. | 82.79% |
| Greenfinch avastrovirus 2 | *Astroviridae* | Greenfinch | *Astroviridae* sp. | 85.37% |
| Passerine avastrovirus 1 | *Astroviridae* | Silvereye | *Astroviridae* sp. | 88.98% |
| Passerine avastrovirus 2 | *Astroviridae* | Greenfinch, Sparrow | *Astroviridae* sp. | 83.16% |
| Passerine avastrovirus 3 | *Astroviridae* | Greenfinch, Sparrow | *Astroviridae* sp. | 83.53% |
| Silvereye avastrovirus 1 | *Astroviridae* | Silvereye | *Astroviridae* sp. | 89.30% |
| Silvereye avastrovirus 2 | *Astroviridae* | Silvereye | *Astroviridae* sp. | 88.10% |
| Sparrow avastrovirus 1 | *Astroviridae* | Sparrow | *Astroviridae* sp. | 84.62% |
| Sparrow avastrovirus 2 | *Astroviridae* | Sparrow | *Astroviridae* sp. | 86.35% |
| Sparrow avastrovirus 3 | *Astroviridae* | Sparrow | *Astroviridae* sp. | 82.79% |
| Sparrow avastrovirus 4 | *Astroviridae* | Sparrow | *Astroviridae* sp. | 83.80% |
| Avian associated calicivirus 1 | *Caliciviridae* | Blackbird, Dunnock, Sparrow | *Avian associated calicivirus 1* | 98% |
| Passerine calicivirus 1 | *Caliciviridae* | Greenfinch, Sparrow | *Caliciviridae* sp. | 69.70% |
| Passerine calicivirus 2 | *Caliciviridae* | Greenfinch, Sparrow | *Caliciviridae* sp. | 60.40% |
| Passerine calicivirus 3 | *Caliciviridae* | Greenfinch, Sparrow | *Caliciviridae* sp. | 66.67% |
| Passerine calicivirus 4 | *Caliciviridae* | Bellbird,  Dunnock, Silvereye | *Avian associated calicivirus 5* | 43.96% |
| Silvereye calicivirus | *Caliciviridae* | Silvereye | *Caliciviridae* sp. | 62.7% |
| Sparrow calicivirus | *Caliciviridae* | Sparrow | *Caliciviridae* sp. | 60.87% |
| Starling calicivirus | *Caliciviridae* | Starling | *Avian associated calicivirus 5* | 59.87% |
| Sparrow deltacoronavirus | *Coronaviridae* | Greenfinch, Sparrow | *Sparrow deltacoronavirus* | 97.75% |
| Goldfinch pegivirus | *Flaviviridae* | Goldfinch | *Leucosticte brandti pegivirus* | 68.29% |
| Greenfinch pegivirus | *Flaviviridae* | Greenfinch | *Leucosticte brandti pegivirus* | 87.07% |
| Passer montanus pegivirus | *Flaviviridae* | Greenfinch | *Passer montanus pegivirus* | 98.61% |
| Passerine pegivirus | *Flaviviridae* | Dunnock,  Greenfinch, Sparrow | *Passer montanus pegivirus* | 44.22% |
| Bellbird genomovirus | *Genomoviridae* | Bellbird | *finch CRESS-DNA virus* | 71.36% |
| Avian associated hepe-like virus 14 | *Hepeviridae* | Bellbird, Silvereye | *Avian associated hepe-like virus 14* | 98.81% |
| Hepatitis E virus | *Hepeviridae* | Greenfinch, Sparrow | Hepatitis E virus |  |
| Passerine hepevirus 1 | *Hepeviridae* | Silvereye | *Hepeviridae* sp. | 41.19% |
| Passerine hepevirus 2 | *Hepeviridae* | Bellbird, Silvereye,  Starling, Sparrow | *Hepeviridae* sp. | 53.13% |
| Silvereye alphaherpesvirus 1 | *Orthoherpesviridae* | Silvereye | [*Gallid alphaherpesvirus 2*](https://www.ncbi.nlm.nih.gov/Taxonomy/Browser/wwwtax.cgi?id=10390) | 65.65% |
| Silvereye alphaherpesvirus 2 | *Orthoherpesviridae* | Silvereye | [*Herpesvirus eupittula14478*](https://www.ncbi.nlm.nih.gov/Taxonomy/Browser/wwwtax.cgi?id=3116695) | 50.75% |
| Avian paramyxovirus 8 | *Paramyxoviridae* | Silvereye | *Avian paramyxovirus 8* | 95.2% |
| Silvereye paraavulavirus | *Paramyxoviridae* | Silvereye | *Paraavulavirus wisconsinense* | 57.06% |
| Silvereye chapahamaparvovirus | *Parvoviridae* | Silvereye | *Fringilla montifringilla Chaphamaparvovirus* | 61.90% |
| Silvereye dependoparvovirus 1 | *Parvoviridae* | Silvereye | *Dependoparvovirus* sp. | 81.25% |
| Silvereye dependoparvovirus 2 | *Parvoviridae* | Silvereye | *Dependoparvovirus* sp. | 57.55% |
| Blackbird anativirus | *Picornaviridae* | Blackbird | *Myna hepatovirus* | 62.30% |
| Chaffinch picornavirus | *Picornaviridae* | Chaffinch | *Avian associated picorna-like virus 22* | 63.45% |
| Dunnock picornavirus | *Picornaviridae* | Dunnock | *Robin hepatovirus 2* | 59.44% |
| Greenfinch hepatovirus | *Picornaviridae* | Greenfinch | *Myna hepatovirus* | 59.02% |
| Greenfinch picornavirus | *Picornaviridae* | Greenfinch | *Avian associated picorna-like virus 34* | 68.57% |
| Passerine picornavirus 1 | *Picornaviridae* | Dunnock, Greenfinch | *Avian associated picorna-like virus 18* | 69.83% |
| Passerine picornavirus 2 | *Picornaviridae* | Greenfinch, Sparrow | *Picornaviridae* sp. | 45.43% |
| Passerine picornavirus 5 | *Picornaviridae* | Silvereye, Starling | *Picornaviridae* sp. | 42.07% |
| Passerine picornavirus 3 | *Picornaviridae* | Silvereye, Starling | *Picornaviridae* sp. | 48.38% |
| Passerine picornavirus 4 | *Picornaviridae* | Bellbird, Silvereye | *Picornavirales* sp. | 85.30% |
| Passerivirus GPS | *Picornaviridae* | Greenfinch, Starling | *Passerivirus GPS* | 93.92% |
| Picornaviridae sp. | *Picornaviridae* | Dunnock | *Picornaviridae sp.* | 95.56% |
| Silvereye hepatovirus | *Picornaviridae* | Silvereye | *Robin hepatovirus 2* | 60.02% |
| Silvereye picornavirus 1 | *Picornaviridae* | Silvereye | *Picornaviridae* sp. | 49.57% |
| Silvereye picornavirus 2 | *Picornaviridae* | Silvereye | *Avian associated picorna-like virus 18* | 34.84% |
| Silvereye picornavirus 3 | *Picornaviridae* | Silvereye | *Petroica picornavirus 1* | 57.81% |
| Sparrow picornavirus | *Picornaviridae* | Sparrow | *Avian associated picorna-like virus 34* | *49.17%* |
| Passerine rotavirus A | *Sedoreoviridae* | Greenfinch, Sparrow | *Rotavirus A* | 85.43% |
| Rotavirus A | *Sedoreoviridae* | Silvereye | *Rotavirus A* | 99.08% |
| Avian orthoreovirus | *Spinareoviridae* | Blackbird, Silvereye,  Sparrow | *Avian orthoreovirus* | 98.73% |
